# Nanomorphological and mechanical reconstruction of mesenchymal stem cells during early apoptosis detected by atomic force microscopy

**DOI:** 10.1101/780759

**Authors:** Xuelian Su, Jizeng Wang, Guangjie Bao, Haijing Zhou, Lin Liu, Qian Zheng, Manli Guo, Jinting Zhang

**Affiliations:** Key Lab of Stomatology of State Ethnic Affairs Commission, Northwest Minzu University, Lanzhou 730030, China; Key Laboratory of Mechanics on Disaster and Environment in Western China, The Ministry of Education of China, Lanzhou University, Lanzhou 730000, China; Key Lab of Oral Diseases of Gansu, Northwest Minzu University, Lanzhou 730030, China

**Keywords:** biomechanics, nanomorphology, apoptosis, bone marrow mesenchymal stem cells, atomic force microscopy, macromolecular crowding

## Abstract

Stem cell apoptosis exists widely in embryonic development, tissue regeneration, repair, aging and pathophysiology of disease. The molecular mechanism of stem cell apoptosis has been extensively investigated. However, alterations in biomechanics and nanomorphology have rarely been studied. Therefore, an apoptosis model was established for bone marrow mesenchymal stem cells (BMSCs) and the reconstruction of the mechanical properties and nanomorphology of the cells were investigated in detail. Atomic force microscopy (AFM), scanning electron microscopy (SEM), laser scanning confocal microscopy (LSCM), flow cytometry and Cell Counting Kit-8 analysis were applied to assess the cellular elasticity modulus, geometry, nanomorphology, cell surface ultrastructure, biological viability and early apoptotic signal (phosphatidylserine, PS). The results indicated that the cellular elastic modulus and volume significantly decreased, whereas the cell surface roughness obviously increased during the first 3 h of cytochalasin B (CB) treatment. Moreover, these alterations preceded the exposure of biological apoptotic signal PS. These findings suggested that cellular mechanical damage is connected with the apoptosis of BMSCs, and the alterations in mechanics and nanomorphology may be a sensitive index to detect alterations in cell viability during apoptosis. The results contribute to a further understanding of the apoptosis from the perspective of cell mechanics.

## 1. Introduction

Stem cells exist extensively in various tissues, such as bone marrow, fat, muscle, hair follicle, dental pulp, and nerves, of organisms. When these tissues are damaged, the body can repair and heal itself. Continuous understanding of stem cells has revealed that these cells play an important role in tissue repair and regeneration, multidirectional differentiation, embryonic development and pathophysiology of disease. [Wu et al., 2007; Sasaki et al., 2008; Amoh et al. 2009]. When large numbers of stem cells are damaged, the body’s ability to repair itself is weakened or lost. For example, chemotherapy agents target rapidly proliferating cancer cells while also inhibit fast-growing stem cells in patients. Those suppressed cells fail to repair the damaged tissue in time, leading to the side effects of chemotherapy, such as alopecia, myelosuppression, immunosuppression and inflammation [Desotelle et al., 2010; Demaria et al., 2017]. In addition, increasing evidence has demonstrated that bone marrow mesenchymal stem cell (BMSC) transplantation and tissue engineering are promising therapeutic strategies for acute brain damage [Tang et al., 2015], spinal cord injury [Rosner et al., 2012], heart injury [Gao et al., 2017] and bone and cartilage defects [Liu et al., 2010, Dahlin et al., 2014]. BMSCs were recruited to the ischemic heart and repaired the injured myocardium after heart infarction [Abbott et al., 2004]. Newly generated cells derived from BMSCs could replace dead cells in injured hearts and vessels. However, BMSC transplantation and engineered tissue construction are hampered by poor survival of stem cells in the recipient sites [Wang et al., 2013; Li et al., 2017]. Apoptosis has become a bottleneck for the further development of stem cell transplantation and tissue engineering. Therefore, understanding cell events during apoptosis of stem cell is imperative.

The apoptosis of stem cells plays an important role in various physiological activities, including embryonic development, homeostasis in normal tissues and aging [Brill et al., 1999; Tower, 2015]. However, abnormal apoptosis of stem cells can lead to disease and/or poor self-repairing capability of the body. One study found that radiation induced apoptosis of BMSCs through mir-22-exotrophic stress, which affected the repair and regeneration of bone tissue in irradiated sites [Liu et al., 2018]. Moreover, the major metabolite 2,5-hexanedione of the organic solvent n-hexane, which is widely used in adhesive preparation, paints, varnishes, oil-based paint, and the food industry, can induce apoptosis of BMSCs via the mitochondria-mediated caspase-3 pathway [Sun et al., 2018; Chen et al., 2015] and nervous lesions [Cui et al., 2007]. Some chemotherapy drugs, such as arsenic trioxide, which significantly inhibited acute leukemia cells, induced mesenchymal stem cell (MSC) apoptosis through the mitochondria-mediated caspase-3 pathway in the patient’s body [Cai et al., 2010]. BMSCs were also significantly damaged by chemotherapy agents in colorectal cancer [Cao et al., 2018]. Therefore, an in-depth understanding of stem cell apoptosis is of great significance for recognizing disease occurrence, drug toxicity, the ability to repair itself, stem cell transplantation and tissue engineering research. In recent years, the molecular mechanism of apoptosis in BMSCs [Chen et al., 2015; Cui et al., 2007; Cai et al., 2010] and malignant cells [Li and Yuan, 2018; Jeong et al., 2018] has been extensively studied. However, the cellular mechanics and nanomorphology during early apoptosis of stem cells have rarely been studied. Therefore, there is a need to understand the biomechanical events of apoptosis in stem cells at the nanoscale.

Bio-type atomic force microscopy (AFM) is an effective device to directly measure the mechanical properties of biological specimens and to quantify the cell morphological changes at the nanoscale level under physiological conditions in a real-time, label-free manner [Muller and Dufrene, 2008; Stark, 2007; Stewart et al., 2012]. The mechanical properties of cancer cells have been extensively explored by AFM during apoptosis [Zhang et al.,2014; Bai et al., 2017]. Our previous studies about apoptosis in cancer cells recognized that the apoptosis is not only closely connected with biological signal cascades but also related to cellular mechanical remodeling [Zhang et al., 2016; Su et al., 2019]. However, alterations in mechanics and nanomorphology during apoptosis have rarely been studied.

Therefore, the present study selected BMSCs to establish an apoptosis model to investigate the alterations in mechanical properties and nanomorphology during early apoptosis. It’s found that the elastic modulus and volume of the cells significantly decreased whereas the surface roughness obviously increased during the first 3 h of cytochalasin B (CB) treatment. These alterations preceded the exposure of biological apoptotic signal PS. Further observation showed that the F-actin fragments filled the cell, and many filaments passed through the membrane and stayed on the cell surface with the disruption of cytoskeleton. These findings suggested that cellular mechanical damage is connected with the apoptosis of BMSCs.

## 2. Results

### 2.1. Identification of BMSCs

The results showed that the expression was negative for the hematogenous markers CD34 and CD45 (0.43% and 0.74%, respectively) on the surface and strongly positive for the stem cell marker CD44 (99.45%). These results collectively suggest that the cells isolated and subsequently cultured were indeed BMSCs and sufficiently pure to meet the experimental requirements.

### 2.2. CB-induced cell growth inhibition

The growth of BMSCs was gradually inhibited by CB. Moreover, the inhibition showed concentration and time dependence (Figs. 1A and B). When the cells were treated with 15 μg/mL CB for 48 h, the maximal inhibitory rate was more than 40%. Therefore, this concentration was selected as the experimental concentration in subsequent procedures. The results suggested that the higher drug concentration and the longer the treatment time was, the greater toxic effects CB had on the cells. The light micrograph indicated the morphological alterations of cells exposed to CB. The untreated cells well spread (shaped as long spindle or long triangle), and the cell boundaries were clearly visible, and the cell boundaries were clearly visible (Fig. 1C). In contrast, for the treated groups, the cells shrank, rounded up little by little, and even detached from the substrate and floated in the medium. The adherent cells were fewer and fewer (Figs. 1D–F).

**Figure 1.**
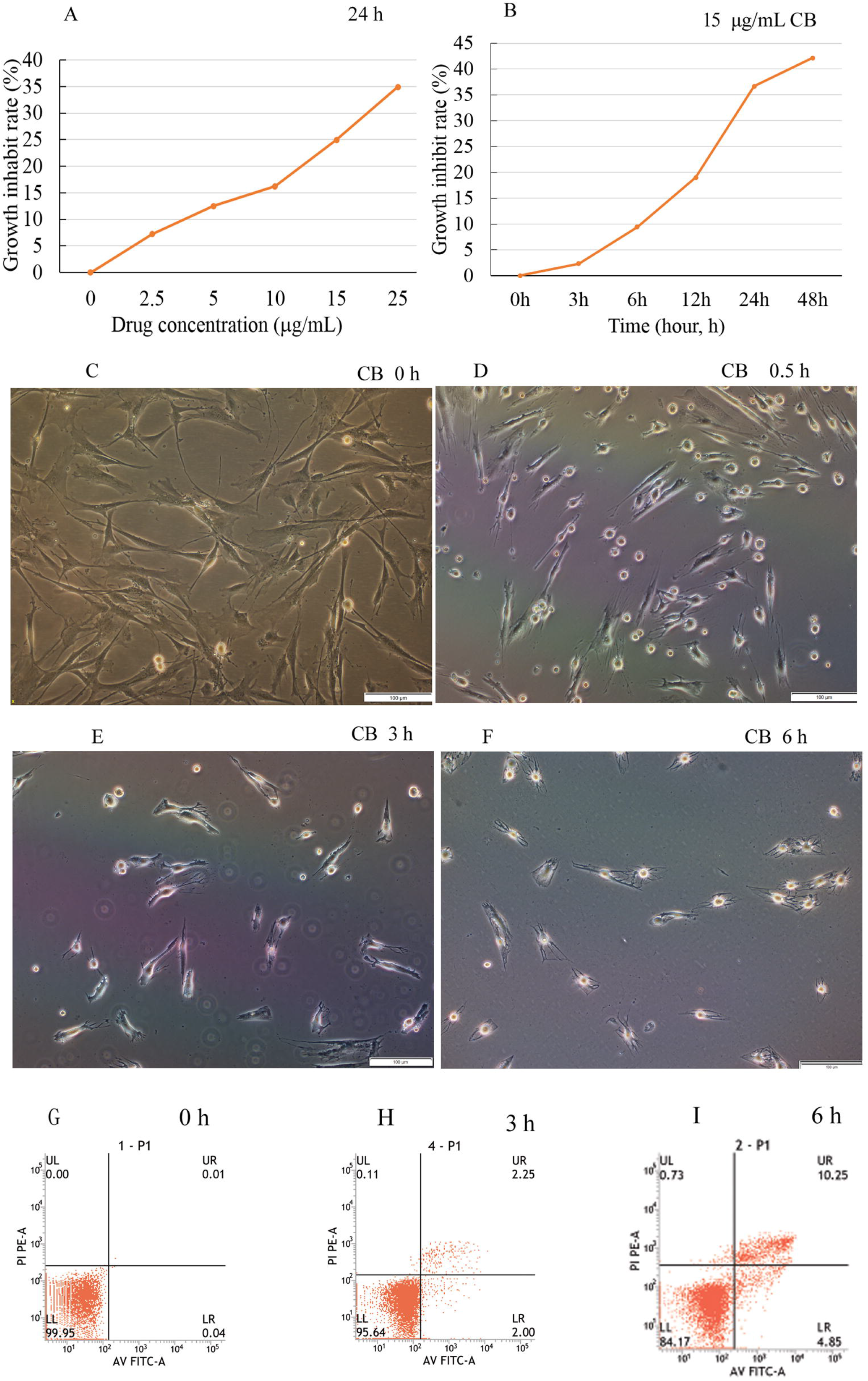
Cell viability and apoptosis assay, morphological changes. The cell death rate increased with the concentrations and treatment time of cytochalasin B, indicating concentration and time dependence (A and B). The light micrograph revealed that the normal BMSCs shaped as long spindle or long triangle, and the boundaries were clearly visible. In contrast, the treated cells shrank little by little, became round, some of them detached from the substrate. The cell boundaries became irregular and the adherent cells were fewer and fewer (C–F). A few apoptotic cells were detected during the first 3 h of treatment with CB (G and H), then the apoptosis rate gradually increased, reaching 15.1% at 6 h (I). Scale bars: 100 μm.

### 2.3. Phosphatidylserine externalized on the membrane

To confirm the biological toxic effects of CB on BMSCs, apoptosis was assessed by FCM using annexin V-FITC/PI double staining. A few apoptotic cells were detected in initial 3 h compared to the control (Figs. 1G and H), and then the rate of apoptosis gradually increased to 15.1% at 6 h (Fig. 2I). The results were consistent with the cell growth inhibition rate and indicated that CB could cause the BMSC apoptosis.

**Figure 2.**
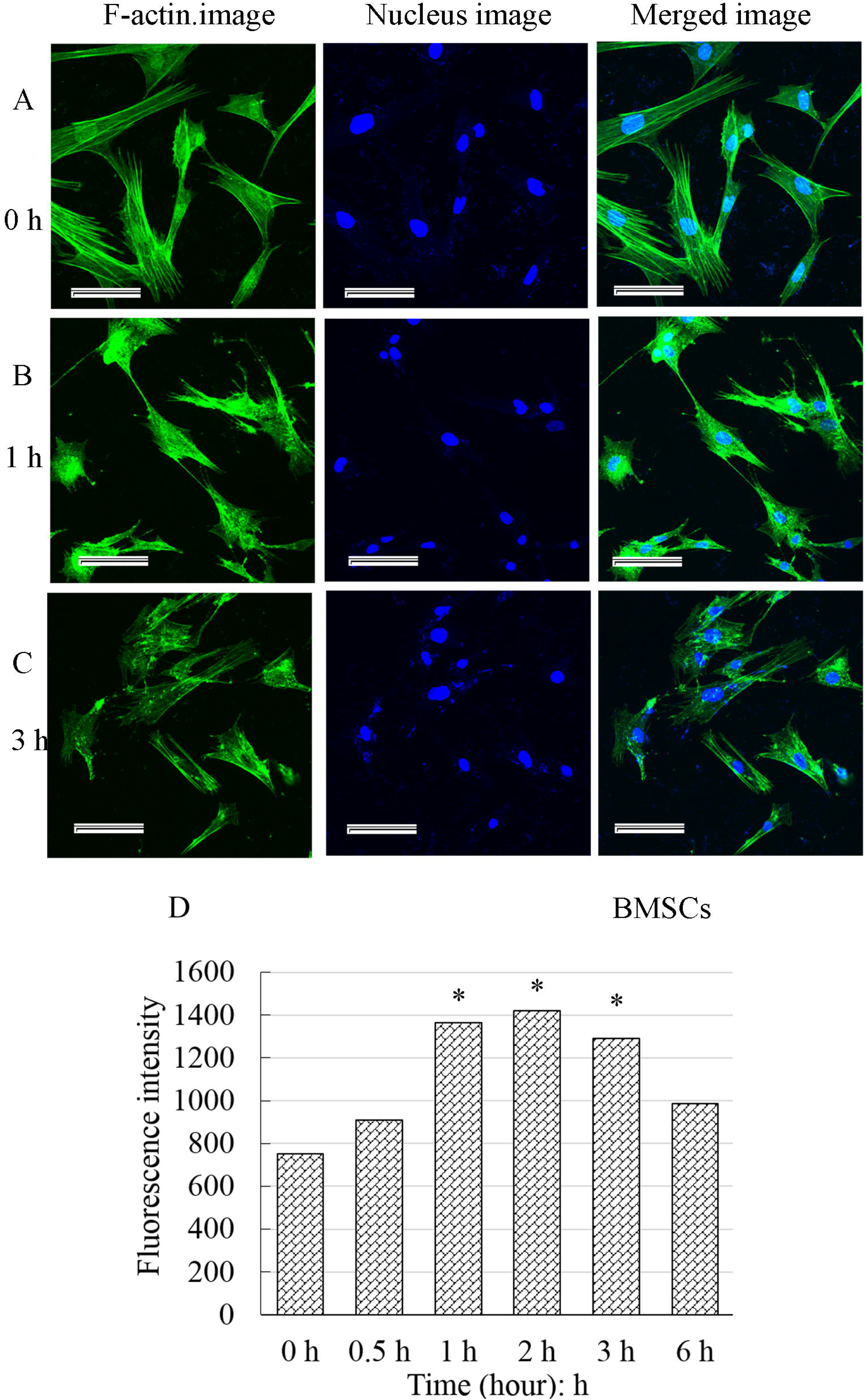
Effect of cytochalasin B on the F-actin cytoskeleton and fluorescence intensity analysis. F-actin uniformly distributed in the cytoplasm below the membrane. The F-actin fibers clustered into bundles (bright green fluorescence) (A). Whereas for the treated cells, the fiber bundles slowly became thinner and smaller, even disappeared and were replaced by smeared or punctate fluorescence fragments (B and C). The fluorescence intensity of F-actin significantly increased, reaching a maximum about 2 h, and then gradually decreased. The results were recorded as the mean ± standard deviation and analyzed by one-way analysis of variance. * *P* < 0.05. Scale bars: 50 μm.

### 2.4. The actin cytoskeleton was disrupted by CB

The F-actin uniformly distributed in the cytoplasm below the membrane and the F-actin fibers clustered into bundles (bright green fluorescence) (Figs. 2A) in control cells. For the treated cells, the F-actin was gradually depolymerized, and the green fiber bundles slowly became thinner and smaller, even disappeared and were replaced by smeared or punctate fluorescence fragments (Figs. 2B and C). The fluorescence intensity of F-actin significantly enhanced, reaching a maximum about 2 h and then increasingly weakened (Fig. 2D). All these alterations revealed that the F-actin cytoskeleton was gradually interrupted by CB, and a large number of actin fragments accumulated in the cells.

### 2.5. Cell surface roughness increased and geometry reconstructed

In the topographic images obtained by AFM, the bright area was the elevated part of the cell, where the nucleus was (Fig. 3). The cell height, diameter, and surface roughness were acquired by cross-sectional analysis of the height-measurement images. The cells in the control group well extend, and their surface was smooth. The texture of the actin bundles was clearly visible (Fig. 3B, 0 h). By contrast, the surface of treated cells became increasingly rough, the periphery of the cells became irregular, and the area of cell extension gradually decreased. The texture of actin bundles disappeared (Figs. 3B). The height of the cell gradually increased (Fig. 3D). In addition, the control cells displayed the smallest roughness (Ra: 667 ± 36 nm, Rq: 677 ± 24 nm). The Ra and Rq values of the treated cells were significantly higher (*P* < 0.05) than those of the control (Fig. 4A and B). These findings strongly confirmed that the surface nanomorphology significantly changed after the cells were treated by CB, exhibiting an obvious time dependence. The height of the cells appeared to increase at first and then slightly decrease, but was always higher than that of the control (Fig. 4C). The cell volume continued to decline and was significantly decreased at 1 h (Fig. 4D). The results suggested that the morphology of cells at nanoscale significantly changed with the continuous depolymerization of F-actin and the collapse of the microfilament cytoskeleton.

**Figure 3.**
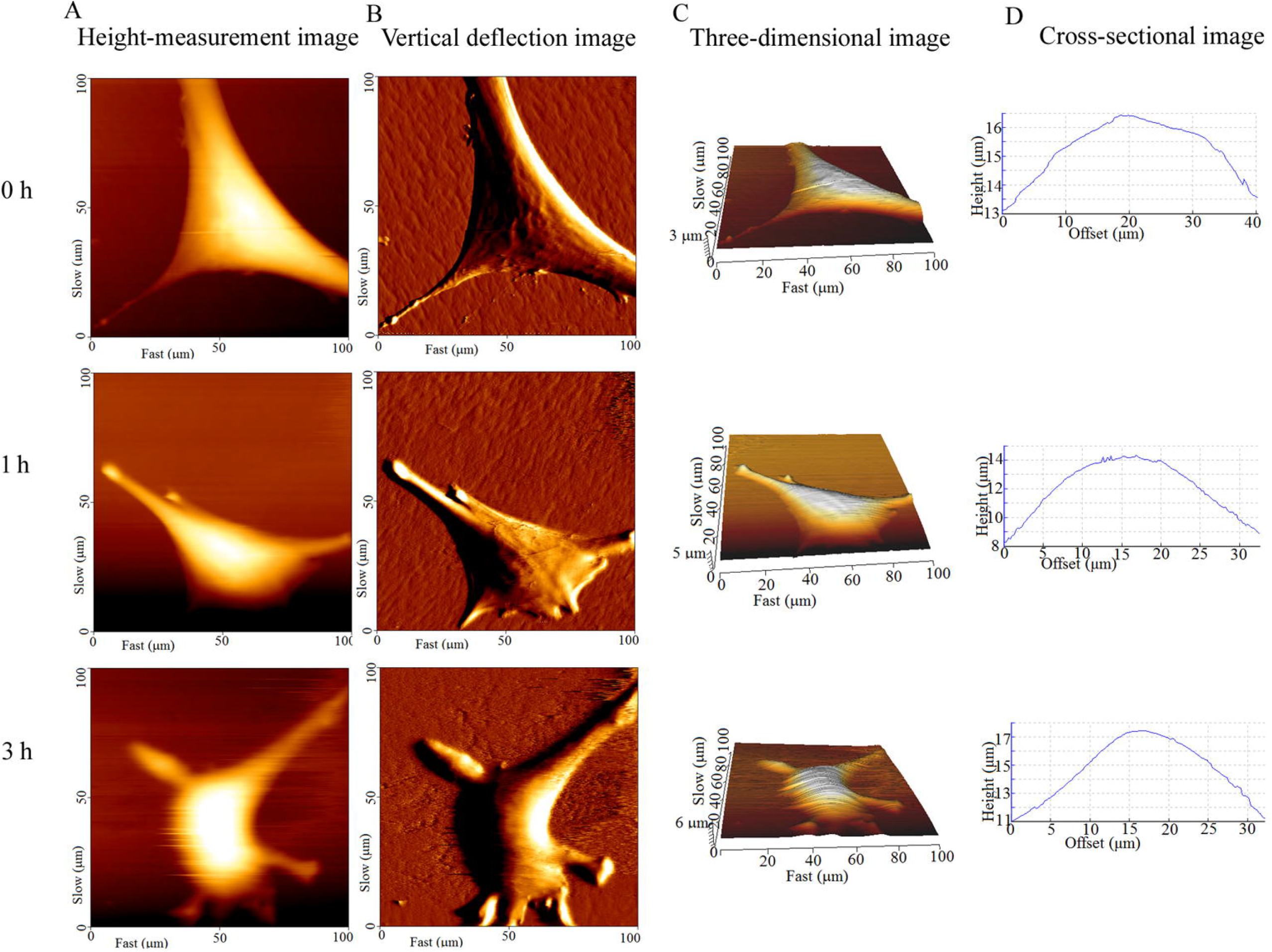
Surface topography of BMSCs captured by AFM at different times. Columns A–D indicated the height-measurement images, vertical deflection images, three-dimensional images and cross-sectional images, respectively. The bright area was the elevated part of the cell, where the nucleus was (A and C). The untreated cells well extend, and their surface was smooth. The texture of the F-actin bundles is clearly visible (B, 0 h). The surface of treated cells became increasingly rough, the periphery of the cells became irregular, and the area of cell extension gradually decreased (A and B, 1 h, 3 h).

**Figure 4.**
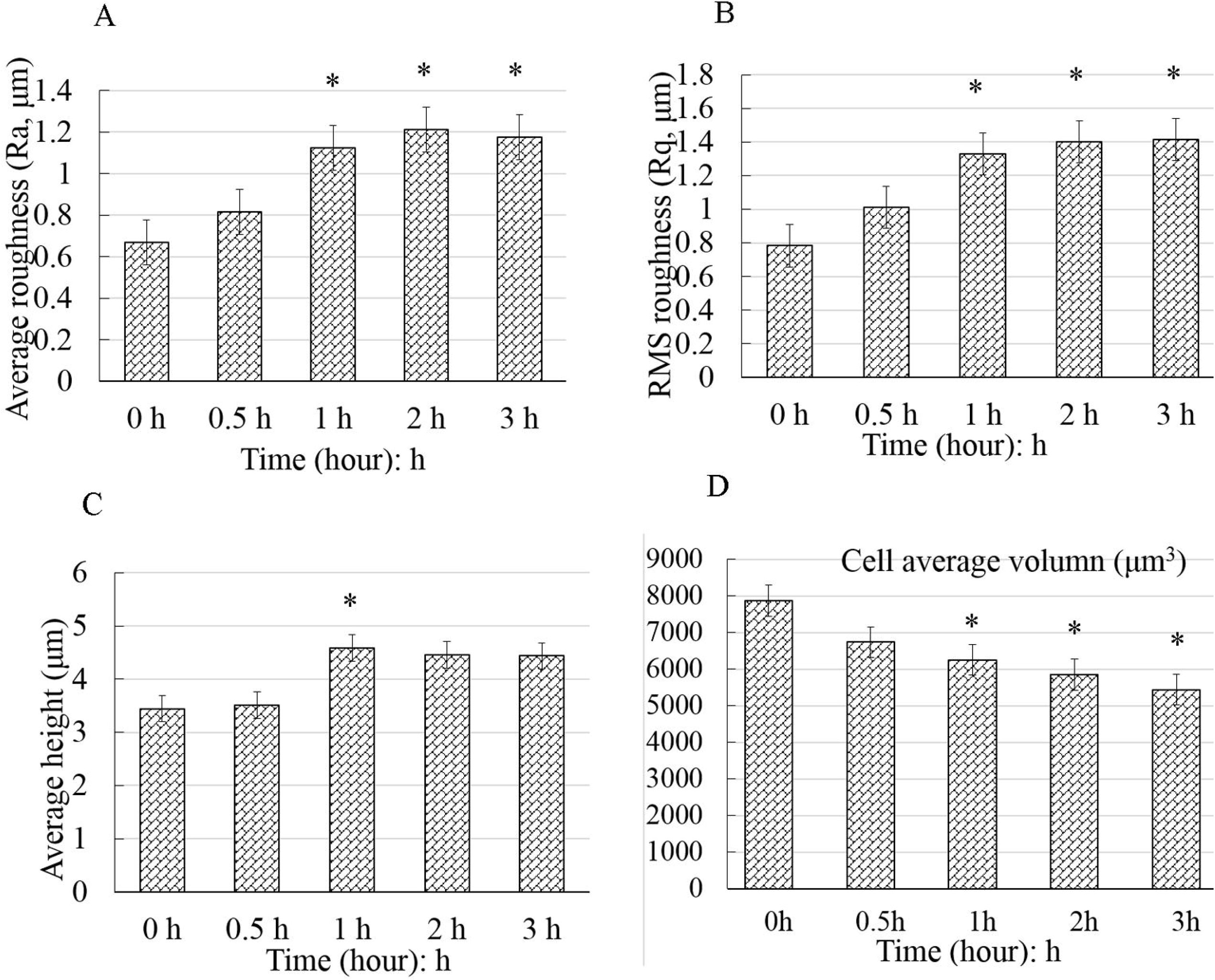
Geometrical reconstruction after the BMSCs were exposed to cytochalasin B. The control cells displayed smallest roughness (A and B). The Ra and Rq values of the treated cells were significantly higher than those of the control cells (*P* < 0.05). The cell height increased at first and then slightly decreased. However, the height of the treated cells was significantly higher than that of the control cells (C). The cell volume continued to decline and significantly decreased at 1 h (D). The data were all displayed as the mean ± S.D. and were analyzed by one-way analysis of variance. * *P* < 0.05. Ra: Average roughness; Rq: Root mean square roughness.

### 2.6. The cell elastic modulus declined

After the height-measurement image was obtained, 10–15 dots/cell were chosen to performed the nanoindentation experiments (Fig. 5A), and the force-distance curves were acquired (Figs. 5C–E). The blue and red lines denoted the approach curves and retract curves, respectively. The elastic modulus *E* of cells were quantified by analyzing the approach curves with Hertz model. Most of the *E* values were distributed around the average value of each group. The average *E* value was 4.55 ± 0.88 MPa for the control cells. After the cells were exposed to CB, *E* continuously decreased to 2.84 ± 0.33 MPa at 3 h (Fig. 5B), indicating that the cells became increasingly softer and softer during the apoptosis induced by CB.

**Figure 5.**
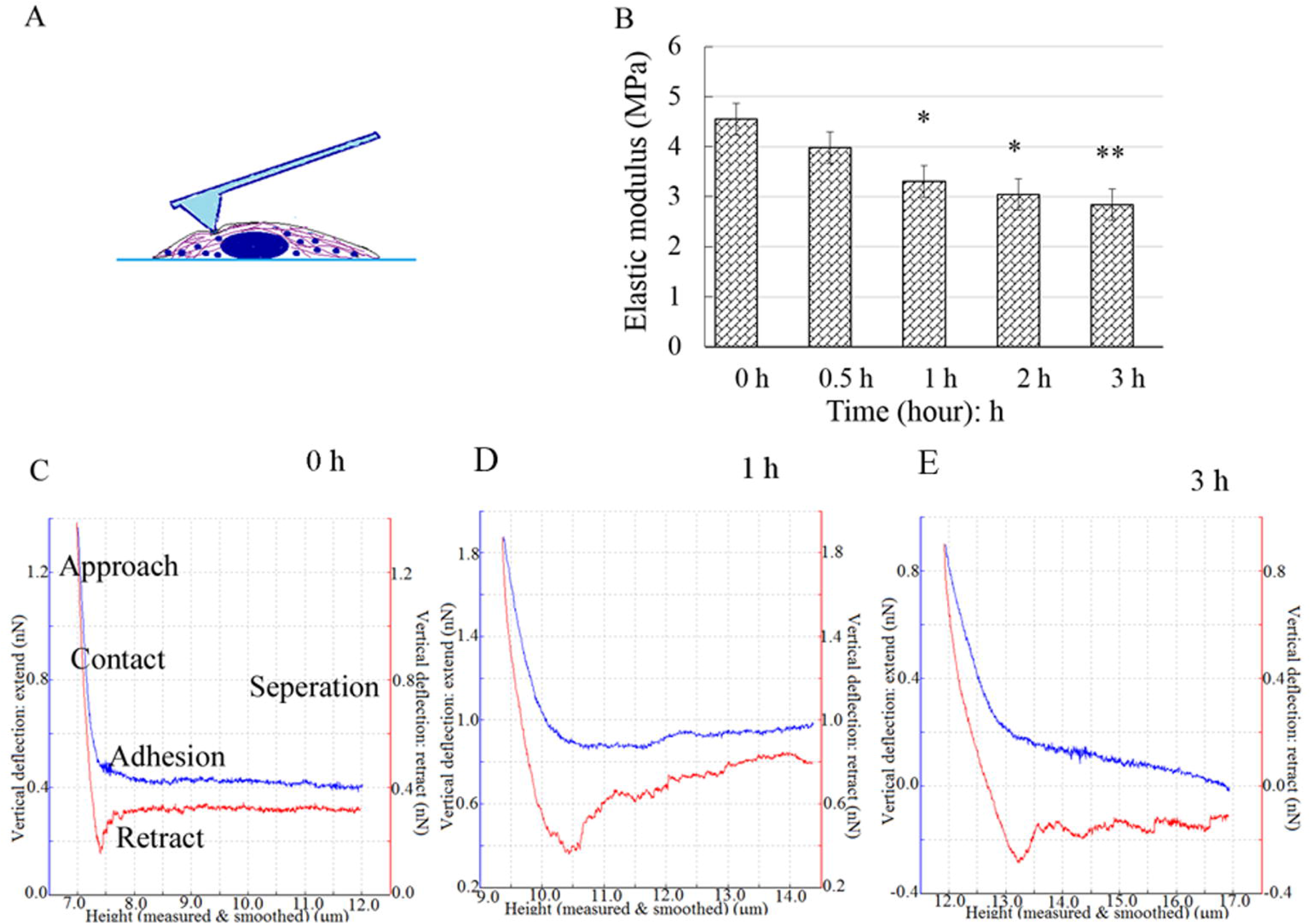
The nanoindentation experiment, force spectra and elastic modulus analysis. A schematic of the indentation experiment: the points were selected a region rich in cytoplasm around the nucleus (A). The force curves were acquired by nanoindentation with AFM (C–E). The blue and red lines represented the approach curves and the retract curves, respectively. The elastic modulus *E* continuously declined and was significantly lower than that of the control at 1 h (B). The results were displayed as the mean ± S.D. and were analyzed by one-way analysis of variance. * *P* < 0.05, ** *P* < 0.01.

The elastic modulus of a cell is mainly depended on the microfilament framework of the cytoskeleton. The mechanical properties of cells are regarded as indicators of cellular biological processes, such as malignant phenotypes, differentiation, and mitosis. Our findings indicated that the elastic modulus and volume significantly decreased, the cell surface roughness obviously increased in initial 3 h of the CB treatment. By comparison, the PS which is a biologically early apoptosis marker, was still undetectable at that time. Therefore, the alterations in cell mechanical properties and nanomorphology occurred more rapidly than the PS exposure, implying that the reconstruction in biomechanics and nanomorphology occurred in early apoptosis.

### 2.7. The surface ultrastructural changed

The surface was smooth for the control group, and only a small number of dots were observed. The pericellular pseudopodia were clearly visible (Figs. 6A and D). In contrast, the cell surface of the treated groups became noticeably rough. A number of curved filaments passed through the membrane and stayed on the cell surface (Figs. 6B, C, E, F). These filaments may be responsible for the increased roughness of cell surface during the BMSC apoptosis. The filaments passed through the membrane could be observed, however, whether these filaments were the broken F-actin fragments disrupted by CB could not be determined.

**Figure 6.**
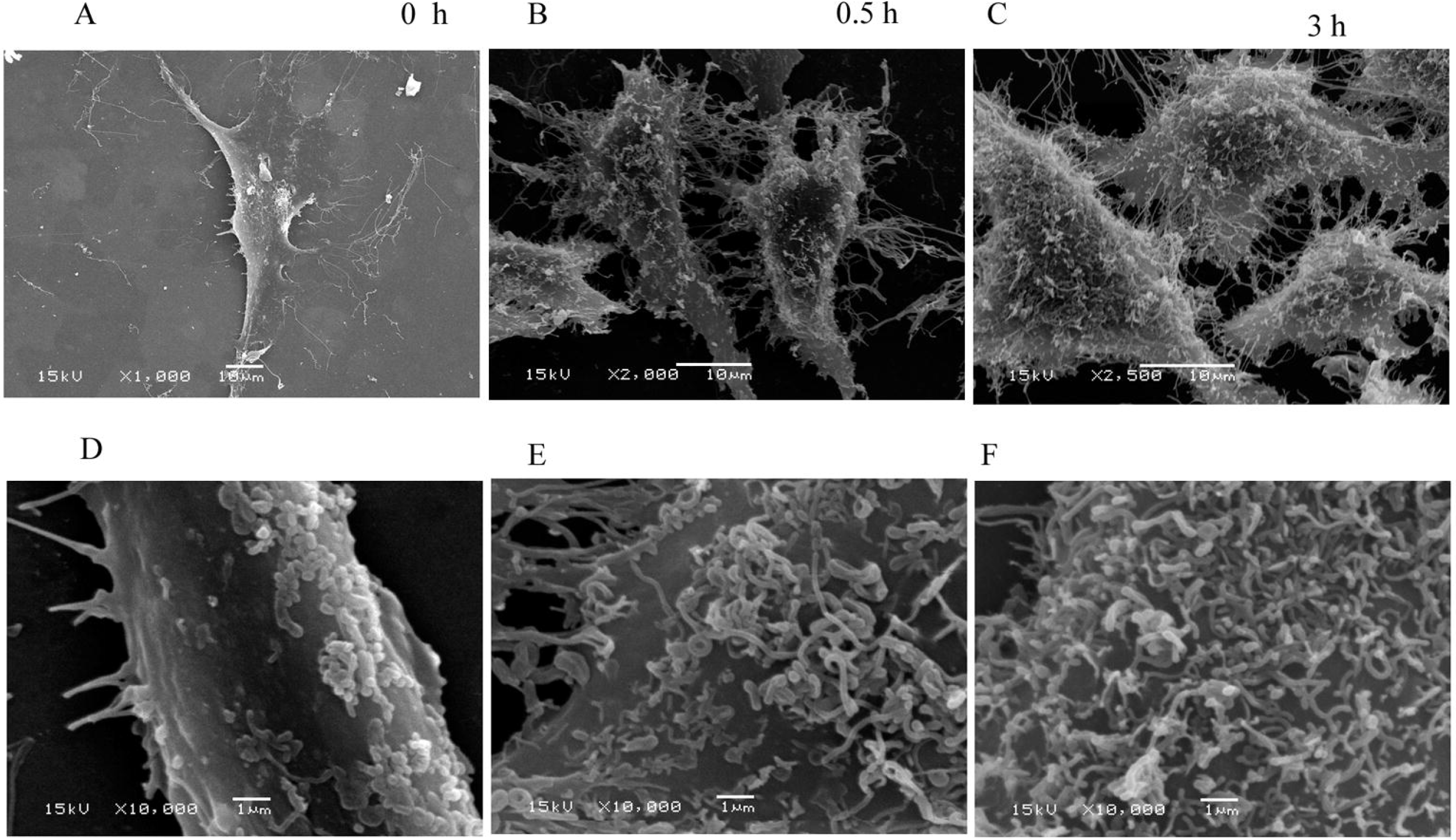
The ultrastructural changes observed by scanning electron microscopy. Overall views of the BMSCs at low magnification (A–C); partial views at high magnification (D–F). The lamellipodia and filopodia were clearly visible around the control cells and a small number of particles were observed on the cell surface (A and D). By contrast, increasingly filamentous structures passed through the membrane and stayed on the cell surface during the cytochalasin B treatment (B, C, E, F).

## 3. Discussion

In the present study, an apoptosis model of BMSCs induced by CB was established. The mechanical events and morphological changes at nanoscale were systematically investigated by AFM and SEM. In early apoptosis, the elastic modulus and volume of the BMSCs significantly decreased while the roughness of the cell surface gradually increased, and numerous filamentous structures crossed the membrane and remained on the surface. These findings are basically consistent with the results of the stiffness, volume and surface roughness during the apoptosis of cancer cells induced by different factors in the literatures [Zhang et al., 2016; Antico et al., 2013]. However, these studies did not delve into why the surface of the cells became rougher. Only our recent study investigated and carefully analyzed the causes of the increased surface roughness of cancer cells for the first time [Su et al., 2019].

The mechanical properties of a cell have been regarded as unique indicators that can directly reflect changes in the vitality state. In particular, alterations in elastic modulus or stiffness have been used as biological markers for cellular phenotypic events and diseases [Haghparast et al., 2013]. In this study, the decrease in elastic modulus and volume suggested that although no obvious biological apoptosis was observed, the cell mechanical vitality was significantly altered within the first 3 h of treatment with CB. Studies have found that microcantilever fluctuations can quantify cell viability and its changes in real time [Kim et al., 2011; Ndieyira et al., 2014]. Therefore, the decline in elastic modulus and volume detected by the AFM microcantilever can reveal decrease in cell viability in our study. Nikolaev and his colleagues investigated the elastic modulus among the viable, impaired membrane and dead cells labeled with annexin V/PI and found that there were significant differences in elastic modulus of these cells [Nikolaev et al., 2014]. In contrast, our findings revealed that CB-treated viable cells that cannot be labeled by annexin V/PI had underwent significant alterations in mechanics, nanomorphology, and geometry. These results suggested that the vitality of these viable cells have changed, and the continuous decrease in cell vitality eventually led to apoptosis. In short, the decrease in elastic modulus and cell volume was consistent with exposure of the early apoptotic signal PS. Moreover, the alterations contribute to further understanding of the mechanical events involved in apoptosis and to monitoring the changes in cell viability before the detection of biological apoptosis signals.

In addition, the cell surface roughness significantly increased with the F-actin cytoskeleton being disrupted by CB. As we know, cytoplasmic membrane blebbing can change the cell surface roughness during apoptosis. However, the membrane blebbing is mediated by caspase-triggered activation of the Rho effector protein ROCK I [Coleman et al., 2001]. Caspases identified as the initiators and effectors or executioners of apoptosis are the products of middle and late apoptosis. Therefore, membrane blebbing should be the phenomenon of late apoptosis, rather than the cause of increased surface roughness in early apoptosis. Vijayarathna et al reported that membrane blebbing was observed by SEM at about 12 h after apoptosis induction [Vijayarathna et al., 2017]. This finding is basically consistent with the results that the caspases were obviously activated at approximately 16 h after the HeLa cells were treated by CB [Kulms et al., 2002; Hwang et al., 2013]. However, in the present study, the cell surface roughness significantly increased within 3 h after CB treatment. The ultrastructural morphology observed (Fig. 6) was also different from that of the membrane blebbing observed by Vijayarathna et al [Vijayarathna et al., 2017]. In fact, fluorescence staining and its intensity analysis indicated that the F-actin cytoskeleton disrupted rapidly, and the actin fragments filled the cells after the BMSCs were treated by CB (Fig. 2). As known, the eukaryotic cells are remarkably crowded with various biological macromolecules [Ellis, 2001]. The F-actin fragments aggravated the degree of molecular crowding in the BMSC. According to Monte Carlo simulation, in a semipermeable biofilm vesicle, larger molecules tend to be redistributed to peripheral region at near the boundary of the vesicle, and some of them protrude across the wall due to the entropic compressing force, as the molecules become more crowded [Shew et al., 2014]. Therefore, the increase in cell surface roughness might arise from the larger molecules protruding across the membrane in extremely crowded conditions, especially F-actin fragments.

In summary, the above data suggested that the cell mechanical alterations and morphological changes at nanoscale are mechanical events that occurred in early apoptosis of BMSCs. In the process of apoptosis, the cell elastic modulus and volume significantly decreased, and the cell surface roughness obviously increased; subsequently, the early apoptosis marker PS was detected, eventually the cells died. In combination with the previous molecular mechanism of stem cell apoptosis [Tower, 2015; Liu et al., 2018; Sun et al., 2018; Chen et al., 2015; Cui, 2007], stem cell apoptosis is not only a biological apoptotic signal cascade process, but is also accompanied by the modification of cell mechanics and nanomorphology. This study is helpful for further understanding and mastering the apoptosis of cells from the perspective of mechanics.

## 4. Materials and methods

### 4.1. Reagents and cell culture

All reagents, including cytochalasin B (CB) and Fetal bovine serum (FBS), were obtained from Merck KGaA (Darmstadt, Germany) unless otherwise specified. An annexin-FITC/ Propidium Iodide (PI) apoptosis detection kit was acquired from MultiSciences (Hangzhou, China). Dulbecco’s modified Eagle’s medium/F-12 (DMEM/F-12) was purchased from HyClone (Utah, USA). Lymphocyte separation medium (LSM) and bovine serum albumin (BSA) were obtained from Solarbio Science & Technology Company.

Isolation, culture and identification of BMSCs: Approximately 3-5 mL bone marrow was extracted from a 3-month-old male goat anesthetized by laughing gas under aseptic conditions and was anticoagulated with 0.2 mL heparin. The marrow was washed with phosphate-buffered saline (PBS, pH 7.2) and centrifuged at 1500 r/min for 10 mins. The supernatant was discarded, and the cell pellets were resuspended in DMEM/F-12 medium supplemented with streptomycin, penicillin and 10% FBS. Then, the suspension was slowly added into the same volume of LSM. After centrifugation at 2200 r/min for 20 min, the bone marrow mononuclear cells were separated and transferred to a new sterile centrifuge tube to be washed three times. The cell pellet was resuspended in DMEM/F-12 medium supplemented with 100 μg/mL streptomycin, 100 U/mL penicillin, 10% FBS, 2 mM/L glutamine, 1% ascorbic acid and 1% nonessential amino acid solution and then plated in 25-cm^2^ culture flasks and cultured in a 37 °C incubator containing 5% CO_2_. The medium was changed every two days. After approximately 7 days of culture, the cells reached 80–90% confluence. The cells were detached by incubation with 0.25% trypsin–EDTA (Invitrogen, USA) and plated at a density of 1×10^4^ cells/cm^2^ for further culture. Cells at passage three were used to identify BMSCs by flow cytometric (FCM, BD FACS Aria, USA) detection of markers, including CD34, CD44, and CD45. BMSCs at passages 3-5 were used for the experiment.

### 4.2. Biological viability and proliferation analysis

The Cell Counting Kit-8 (CCK-8) assay was used to estimate the biological cytotoxicity of the cells according to the manufacturer’s instructions. The BMSCs were plated at 5×10^3^ cells/well in a 96-well plate and incubated overnight. Each group had three replicates (wells). The medium was substituted with fresh medium containing CB at various concentrations (0, 5, 10, 15 and 25 μg/mL) and incubated for 24 h. Then, 0.01 mL assay reagent was added to each well. After 1.5 h, the 96-well plate was analyzed using an enzyme-linked immunosorbent assay. The optical density of living cells was read at 450 nm in a microplate reader (Synergy HT; BioTek Instruments, Inc. Winooski, VT, USA).

### 4.3. Apoptotic analysis of BMSCs by annexin V-FITC/PI

During early apoptosis, the intracellular Ca^2+^ concentration increases, which causes the phosphatidylserines (PS) to translocate from the inner to the outer leaflet of the cell membrane [Suzuki et al., 2010; Martin et al., 1995]. Therefore, the study applied the PS exposure to determine the early apoptosis, and design the subsequent experiments. Annexin has a high affinity to PS and the translocation of PS was identified by an annexin V. The early apoptosis was analyzed by flow cytometry (FCM) through the annexin V-FITC/PI apoptosis detection kit. Approximately 5×10^5^ cells seeded on a 60-mm diameter Petri dish, incubated for 24 h, treated with CB for 0, 3, 6, 12 h and collected by trypsinization. The cell pellets were resuspended in 100 μL binding buffer and stained with 5 μL annexin V-FITC and 5 μL PI for 30 min at room temperature (approximately 25 °C) in the dark. The samples were analyzed with a FACSCalibur flow cytometer (BD Biosciences, Franklin Lakes, NJ, USA).

### 4.4. F-actin cytoskeleton fluorescence staining and visualization

Cytochalasin B is a cytotoxic agent that interferes with the polymerization of actin filaments (F-actin). To visualize the alteration in F-actin framework, the BMSCs were grown on round coverslips (preplaced sterile coverslips in a 24-well cell culture plate) at a density of 4,000 cells/cm^2^, stained with fluorescein isothiocyanate (FITC)-phalloidin. The specimens were rinsed with PBS, fixed with 4% cold paraformaldehyde, permeated with 0.2% Triton-X100 in PBS, blocked with 2% BSA, and incubated with phalloidin for 1 h at room temperature in the dark. The nuclei were labeled with 0.1 mg/ml DAPI. The coverslips were sealed on glass slides with glycerin sealant, observed and photographed with a laser scanning confocal microscope (LSCM, Olympus FV1000, Japan) within a week. The mean fluorescence intensity was analyzed using FV10-ASW 4.1 Viewer software (Olympus Corp., Japan). No any filtering or adjustments were performed.

### 4.5. Single-cell imaging and nanoindentation experiments

A bio-type Nano-Wizard III AFM (JPK Instruments, Germany) was employed to detect the topographical changes and quantify the alterations in the mechanical properties of living cells. The imaging and nanoindentation experiments were performed according to previously described operating procedures [Zhang et al., 2016; Su et al., 2019]. The microscope was combined with an inverted optical microscope (Carl Zeiss, Germany) that was used to select the ideal cell. Silicon nitride cantilevers (PNP-DB, Nano World, Neuchatel, Switzerland) with a spring constant of 0.030 N/m (fo: 17 kHz) were applied. Prior to the measurement, the spring constants of the cantilever were calibrated with JPK Instruments software 4.2.61. A tip with a spring constant of 0.028–0.030 N/m was used in the subsequent experiments. The AFM imaging and nanoindentation experiments were implemented under intermittent mode using a square pyramidal tip with a 25° half-opening angle. When the tip intermittently interacts with the cells, the force-sensitive microcantilever fluctuated in three dimensions. The deflection was captured through the alteration of a laser projected on the cantilever. Approximately 1×10^4^ cells/dish were seeded in 35-mm Petri dishes. The experiments were conducted in cell culture medium at 37 °C using a liquid temperature-controlled chamber. About 10–15 cells were imaged for each group. After topography scanning, 10–15 dots around the nucleus (this area is rich in cytoplasm, avoiding the influence of the substrate) were chosed to acquire force-distance curves by indentation. The interactions between the tip and the sample caused the cantilever to deflect, which was recorded as a function of the relative sample position, that is, a force-distance curve. The elastic modulus *E* (cell stiffness) was calculated through analysis of a series of 130-150 curves with JPK Instruments data processing software.

### 4.6. Quantitative analysis of alterations in nanomorphology and geometry

The height of the cell was defined as the distance between the top and bottom of the cell. The height, diameter and surface roughness were quantified by cross-sectional analysis of the height-measurement images. The average roughness (Ra) and Root-Mean-Square roughness (Rq) are crucial parameters for understanding the surface morphology of living cells at the nanoscale. The roughness provided quantitative data regarding the reconstruction of surface nanomorphology after the F-actin cytoskeleton was disturbed. The cellular volume is also another important indicator for the state of cell viability. The cell was regarded as half an oblate ellipsoid, its volume can be calculated according to the following equation [Hessler et al., 2005]:

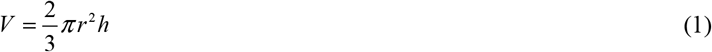

where *V* is the volume, *r* is the radius, and *h* is the height.

### 4.7. Elastic modulus measurement

The height-measurement images and the force-distance curves were obtained by the sensitive cantilever fluctuations. The cell was considered an elastomer of homogeneous structure. Thus, the cell elastic modulus *E* (cell stiffness) was calculated according to the Hertz model and the approach curves [Zuk et al., 2011; Geiger et al., 2009]. The referential equation that gives the relation among indentation force, elastic modulus and depth is as follows:

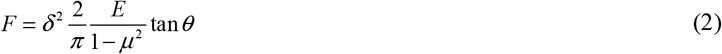

where *E* is the elastic modulus, *F* is the loading force, *μ* is the Poisson’s ratio of the samples, *δ* is the indentation depth, and *θ* is the half-opening angle of the tip.

### 4.8. Cell surface observed by scanning electron microscopy

Prior to obvious biological apoptosis, the surface roughness measured by AFM significantly increased. To further elucidate the phenomenon, the cells were observed by scanning electron microscopy (SEM). Approximately 2-5×10^3^ cells were seeded onto sterile coverslips in a 24-well cell culture plate, cultured for 24 h, and treated by CB. The samples were rinsed with PBS, fixed with precooled glutaraldehyde solution at 4 °C overnight, and made into ultrathin slices. The cells were observed and captured by SEM (JSM6380 LV, Japan).

### 4.9. Statistical analysis

The data were recorded as the mean ± standard deviation (SD) and analyzed using SPSS 22.0 (Statistical Product and Service Solutions, Stanford University, USA). Statistical differences were performed using one-way analysis of variance. *P*-values <0.05 were considered significantly statistical differences. A single asterisk (*) indicates significant difference (*P* < 0.05), and double asterisks (**) denote extremely statistical difference (*P* < 0.01).

## Conclusion

These findings suggested that cellular mechanical damage was connected with the apoptosis of BMSCs, and the reconstructions in mechanics may be a sensitive index to detect alterations in cell viability during apoptosis.

## Ethics approval

The Bioethical Committee of the Northwest Minzu University approved the use of the samples for this research.

## Acknowledgments

The authors thank the Key Lab of Stomatology of State Ethnic Affairs Commission (Northwest Minzu University). We also thank Ms. Fei Tang and Mr. Ming-zhong Chen for their technical assistance with flow cytometry analysis and CLSM imaging, respectively.

## Author contributions

X S performed the majority of the experiments. J W, and G B designed the experiments and coordinated the project. The manuscript was written by X S and revised by J W and G B. H Z, L, Q Z, M G and J Z participated in the in vitro study. All the authors reviewed the manuscript.

